# Exploring ligand binding pathways on proteins using hypersound–accelerated molecular dynamics

**DOI:** 10.1101/2020.04.06.026930

**Authors:** Mitsugu Araki, Shigeyuki Matsumoto, Gert-Jan Bekker, Yuta Isaka, Yukari Sagae, Narutoshi Kamiya, Yasushi Okuno

## Abstract

Capturing the dynamic processes of biomolecular systems in atomistic detail remains difficult despite recent experimental advances. Although molecular dynamics (MD) techniques enable atomic-level observations, simulations of “slow” biomolecular processes (with timescales longer than submilliseconds) are challenging, due to current computer speed limitations. Therefore, we developed a new method to accelerate MD simulations by high-frequency ultrasound perturbation. The binding events between the protein CDK2 and its small-molecule inhibitors were nearly undetectable in 100-ns conventional MD, but the new method successfully accelerated their slow binding rates by up to 10–20 times. The accelerated MD simulations revealed a variety of microscopic kinetic features of the inhibitors on the protein surface, such as the existence of different binding pathways to the active site. Moreover, the simulations allowed estimating the corresponding kinetic parameters and exploring other druggable pockets. This method can thus provide deeper insight into the microscopic interactions controlling biomolecular processes.

## Main text

The microscopic observation of biomolecular processes such as protein folding, protein interactions, and enzyme reactions, most of which occur on timescales ranging from microseconds to seconds (*1*), is of great interest to the molecular biology community. Although molecular dynamics (MD) simulations enable atomic-level observations, they are limited to several microseconds on standard high-performance computers and are thus normally applicable only to relatively fast processes (*2*). Recently, the kinetics of slower protein interaction processes were explored through long MD simulations spanning timescales of tens of microseconds to milliseconds (*3-5*), which were achieved through the development of special-purpose supercomputers for high-speed simulations (*e.g.*, ANTON (*6*)) and/or algorithms to aggregate many short simulations (*e.g.*, Markov state models (MSMs) (*7*)). Unfortunately, MD-specific supercomputers such as ANTON are accessible only to a limited number of researchers due to their high costs. While MSMs have a lower requirement for simulation power, this method is very sensitive to the choice of the hyperparameters (*8*), which makes MSM approaches less than straightforward to use.

To overcome these problems, we have developed a new MD simulation method that utilizes high-frequency ultrasound (hereafter denoted as hypersound) shock waves to accelerate the dynamics. This method falls into the category of nonequilibrium MD simulations under external field perturbation (*9, 10*). Its key advantage is that it allows naturally “slow” processes such as those mentioned above to be frequently and directly captured in a series of single MD trajectories performed on standard high-performance computers. In the experimental field, ultrasound irradiation procedures have been applied to accelerate various kinds of chemical reactions (*11, 12*) and to synthesize nanoparticles (*13*). This acceleration is considered to be induced by acoustic cavitation (*i.e*., the repeated growth and collapse of cavitation bubbles formed by the ultrasound waves, which generate local high-temperature/pressure regions in solution) (*14*). Inspired by these results, in this study, we first analyzed the hypersound-dependent behavior of a liquid water model to test the effect of shock waves with a protein-size wavelength. Next, to assess the effect of hypersound acceleration on biomolecular processes, we performed short (100–200 ns) simulations to capture the slow binding of small-molecule inhibitor compounds (CS3 and CS242) to cyclin-dependent kinase 2 (CDK2) (*15*), as a representative system in which the binding event would be nearly undetectable in standard MD. The simulations showed a significant acceleration of the binding process under hypersound irradiation compared to standard MD simulations. The accelerated simulations revealed the existence of various conformationally and energetically diverse binding pathways, suggesting that the assumption of a single pathway/transition state made in conventional kinetic models may be inaccurate. The present method allowed not only the estimation of kinetic parameters of slow binding inhibitors, but also the full exploration of druggable sites. This approach would thus be helpful for efficiently understanding the microscopic mechanism of slow biomolecular processes.

To simulate hypersound shock waves with protein-size wavelengths, their frequency was set to 625 GHz (corresponding to a period of 1.6 ps) (Fig. S1, Materials and Methods, and Supplementary Material), which is more than 100 times higher frequency than that of currently used ultrasound waves (*16*). The hypersound-perturbed MD simulation of liquid water at 298 K showed the generation and propagation of a high-density region (Fig. 1A and Movie S1). Then, we analyzed the MD trajectory focusing on wave propagation along the X direction as an representative example (Fig. 1B–D). As the X coordinate of the first wave reached 4 nm at a simulation time of 1.7 ps after passing through X = 2 nm at 0.7 ps (Fig. 1D), the propagation speed of the shock wave could be estimated to be 2,000 m/s, which is similar to the speed of sound in water (∼ 1500 m s^-1^). The wavelength of the simulated shock wave was estimated to be 3.2 nm (= 2000 m s^-1^ × 1.6 ps), which corresponds to the hydrodynamic radii of globular proteins consisting of 300–400 residues (*17*); this confirms the successful generation of hypersound waves, appropriate to perturb biomolecular processes. The pressure (*px*) and kinetic energy (*kx*) of the liquid water model also exhibited periodic fluctuations with the same phase as the density, respectively reaching ∼ 2000 atmospheres and 0.4–0.5 kcal/mol (corresponding to 400–500 K) at the center (X = 4 nm) of the simulation box (Figs. 1B, C, and E). In contrast, the macroscopic properties of liquid water were not affected by the hypersound irradiation, except for the diffusion constant, which slightly increased to 6.30 ± 0.10 × 10 ^-5^ cm^2^ s^-1^, equivalent to the corresponding parameter of bulk water at 305 K (Table S1). These results demonstrate that hypersound irradiation of a liquid solvent generates local higher pressure and temperature regions appropriate for promoting chemical processes (*14*) without altering the macroscopic properties of the liquid.

**Figure 1.**
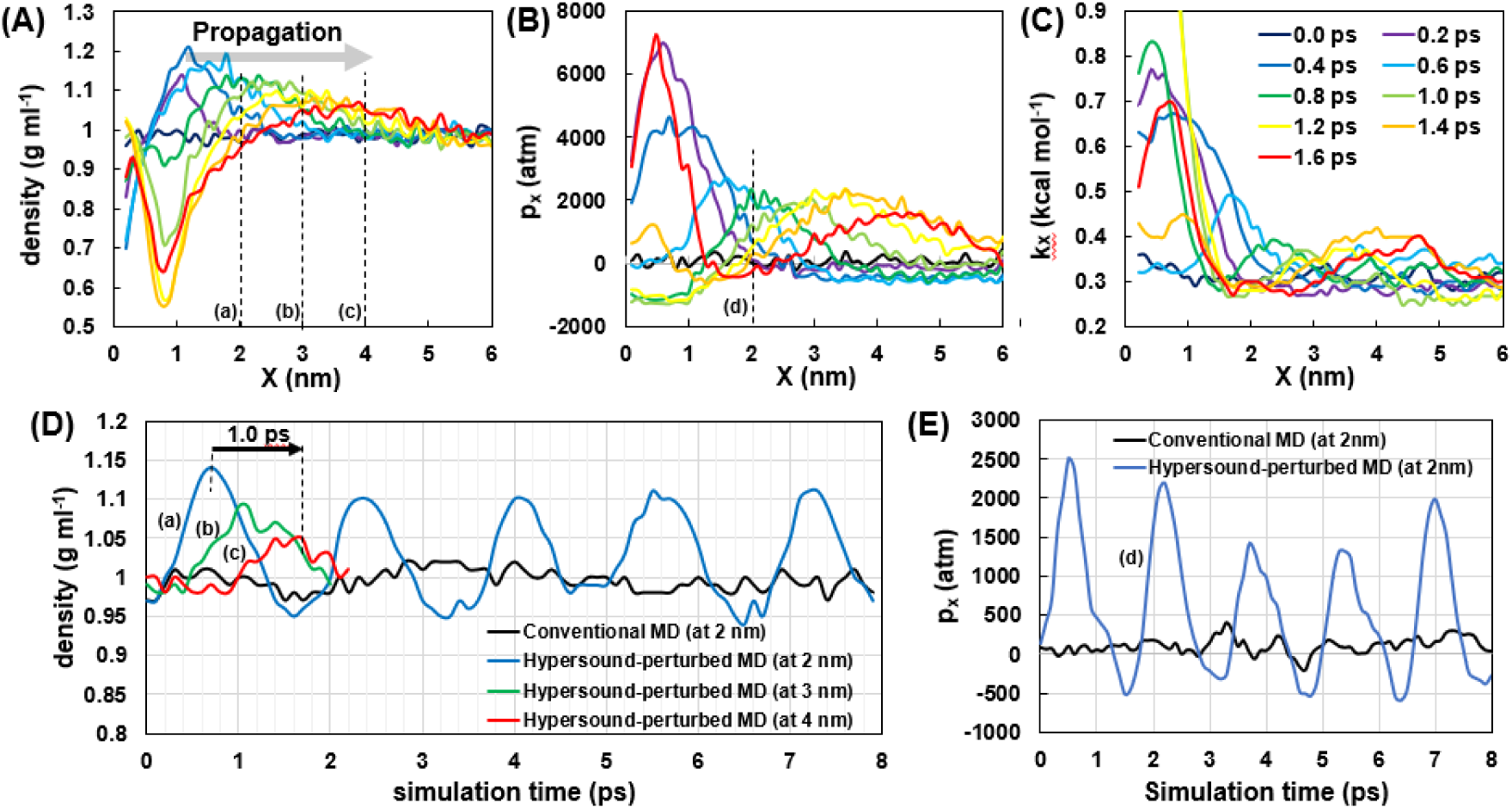
Hypersound-perturbed water dynamics at 298K. (A-C) Spatial variation of (A) mass density, (B) pressure in the +X direction (p*x*), and (C) X component of kinetic energy (*kx*), measured at different simulation times. (D-E) Time dependence of (D) mass density and (E) the pressure, measured at different X positions; the corresponding positions are shown in (A) and (B). Shock waves were generated in the X = 0–1 nm region (X_0_ surface in Fig. S1) of the simulation box. Further details can be found in the Supplementary Material.

We next assessed the influence of the shock wave perturbation on the association between the CDK2 protein and its slow-binding ATP-competitive inhibitors CS3 and CS242. The probability of observing the ligand binding event in conventional MD simulations of 100 ns was estimated to be only 0.7% (= 2/283, corresponding to 2 out of 283 MD runs resulting in binding) for CS3 and 0.5% (= 2/369) for CS242 (Table S2). On the other hand, the CS3 and CS242 binding probabilities observed in 100-ns long hypersound-perturbed MD simulations were 11.9% (= 27/226) and 3.9% (= 14/362), respectively (Table S2), showing that the perturbation successfully increased the association rate by 17.0 times (= 11.9/0.7%) for CS3 and 7.8 times (= 3.9/0.5%) for CS242. MD trajectories that exhibited stable ligand binding were extended to 200 ns to observe the behavior of the bound ligand, and based on their percentages, the association rate constants (*k*_*on*_) for CS3 and CS242 under hypersound irradiation were estimated to be 3.68 × 10^6^ and 1.92 × 10^6^ M^-1^ s^-1^ (see also Materials and Methods section), respectively, which was consistent with the order of their experimental *k*_*on*_ values, measured without any perturbation (*15*) (Table 1). This analysis proved the effectiveness of the hypersound-perturbed simulations in enhancing the sampling of infrequent binding events; this approach can thus be applied to extract further atomic-level information on these processes, as follows.

**Table 1.** Kinetic parameters of the CDK2-ligand binding process determined by conventional and hypersound-perturbed MD simulations (see also Materials and Methods section).

| | Association<br>rate constant<br>$k_{on}$ ($M^{-1} s^{-1}$) | Activation<br>energy<br>$E$ (kcal mol$^{-1}$) | Diffusion<br>constant<br>$D$ ($\times 10^{-5}$ cm$^2$ s$^{-1}$) | Steric factor<br>log($P$) | Frequency<br>factor<br>log($A$ ($M^{-1} s^{-1}$)) | Effective<br>temperature<br>$T$ (K) |
| --- | --- | --- | --- | --- | --- | --- |
| CS3 (conventional MD) | $3.35 \times 10^5$<sup>a</sup> | $4.8 \pm 2.2$ | $1.87 \pm 1.33$ | $-1.05 \pm 1.64$ | $9.10 \pm 1.69$ | 298 |
| CS3 (hypersound MD) | $3.68 \times 10^6$ | $4.8 \pm 2.2$ | $10.62 \pm 6.98$ | $-1.05 \pm 1.64$ | $9.86 \pm 1.68$ | 316 |
| CS242 (conventional MD) | $3.21 \times 10^4$<sup>a</sup> | $8.4 \pm 2.3$ | $1.55 \pm 1.01$ | $0.52 \pm 1.70$ | $10.59 \pm 1.73$ | 298 |
| CS242 (hypersound MD) | $1.92 \times 10^6$ | $8.4 \pm 2.3$ | $9.87 \pm 7.00$ | $0.52 \pm 1.70$ | $11.39 \pm 1.74$ | 357 |

<sup>a</sup> Experimentally determined $k_{on}$ values retrieved from the Community Structure-Activity Resource (CSAR) database (15).

The accelerated MD simulations revealed that multiple transitions among different conformations took place within each individual binding pathway (see Fig. 2A and Movie S2 for CS3 and Fig. 2B and Movie S3 for CS242). This emerges from the inspection of the 27 (CS3) and 14 (CS242) binding pathways observed in the hypersound-perturbed MD simulations, a few representative cases of which are shown in Figs. S2 and S3. It should be noted that these pathways contain those observed in the conventional and existing generalized-ensemble MD simulations (*18*) (Fig. S4). The potential energy trajectories (also displayed in the figures) reveal the occurrence of multiple energy barriers along each binding pathway, and show that the position and height of the highest-energy transition state depend on the binding pathway (Fig. 2C). The trajectories indicate that the ligand tends to adopt energetically unstable configurations upon (i) entry into the CDK2 pocket (Figs. 2A, S2A, and S3A) or (ii) conformational rearrangement in the pocket interior (Figs. 2B, S2B, and S3B); these effects have not been previously captured by ensemble-averaged kinetic experiments (*15*)(*19*). Ligand unbinding was also observed in some of these trajectories, most of which also exhibited different binding and unbinding pathways (Figs. S2C and S3C); this suggests that the conventional kinetic model based on identical binding/unbinding pathways is not always valid at the single-molecule level. The trajectories of individual ligand molecules captured by the hypersound perturbation approach revealed the complex microscopic nature of the CDK2-inhibitor binding kinetics, highlighting the effectiveness of this approach in exposing effects not accessible by other experimental and computational techniques.

**Figure 2.**
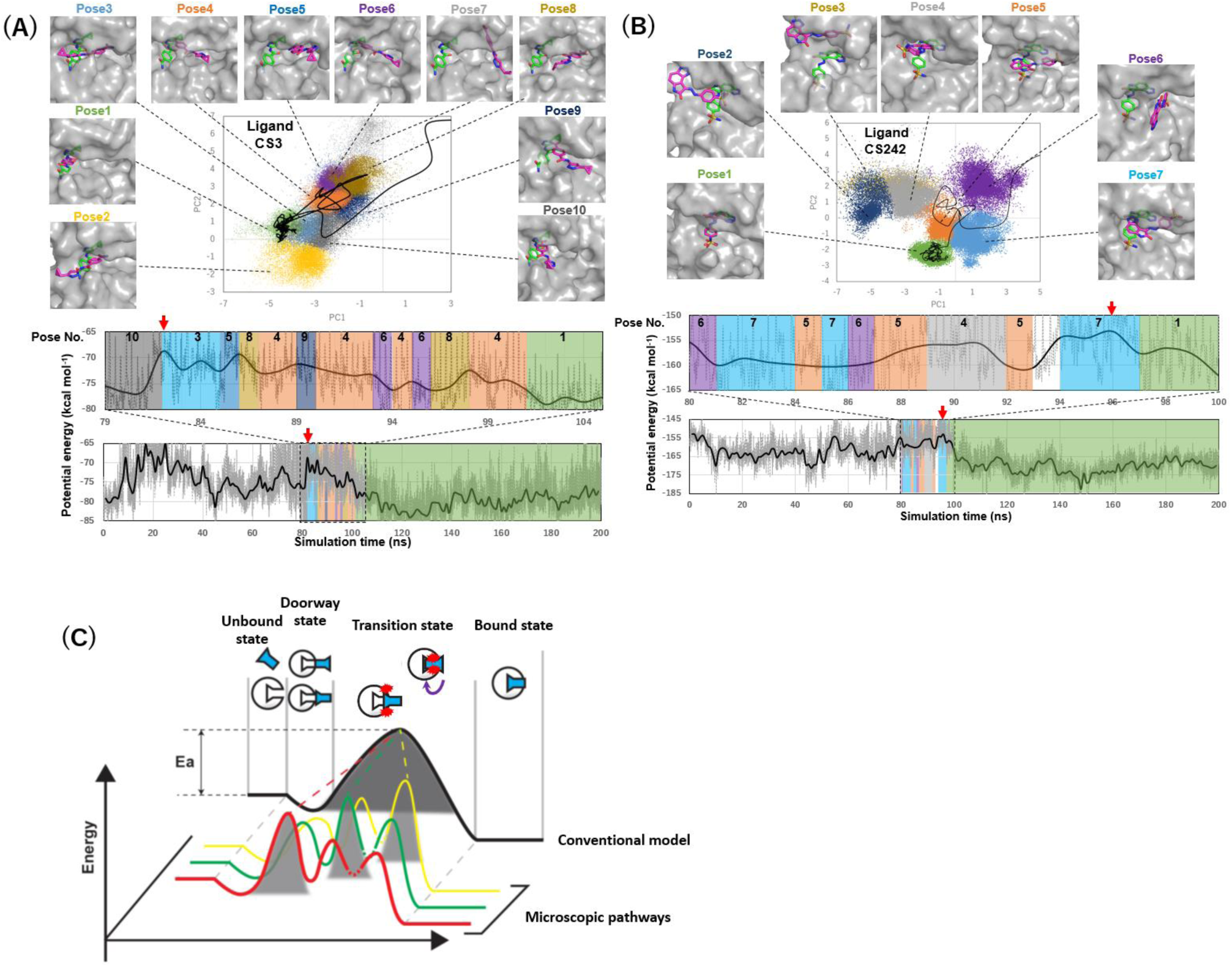
Microscopic binding pathways of CDK2 inhibitors. (A, B) Representative binding pathways of (A) CS3 and (B) CS242 ligands to the ATP-binding pocket of CDK2. (top) Projections of binding conformations observed in the whole set of MD trajectories (colored dots) and of a representative binding pathway (black line) onto the first and second principal components (PC1 and PC2) calculated from principal component analysis (PCA; see Supplementary Material). Ten (CS3) and 7 (CS242) representative binding poses (magenta sticks) on CDK2 (gray surfaces) are shown alongside the crystallographic pose (green sticks), the closest conformation to which was assigned as Pose 1. (bottom) Potential energy trajectory corresponding to the pathway shown in the PCA map. The highest-energy transition state is indicated by a red arrow. The upper panel shows an enlarged view of the potential energy trajectory close to the highest-energy transition state. Note that transition states occur (A) immediately before/after the ligand enters the CDK2 pocket and (B) during conformational rearrangements taking place after pocket entry. (C) Schematic illustration of microscopic and macroscopic kinetic models. The conventional kinetic model assumes a single binding pathway with a single transition state. However, at the single-molecule level, the ligand binds to the protein through multiple pathways with different highest-energy transition state conformations.

By averaging the energy barriers observed in 12 (CS3) and 7 (CS242) trajectories that exhibited stable ligand binding (Table S2), the activation energies for CS3 and CS242 binding to CDK2 were estimated to be 4.8 and 8.4 kcal/mol, respectively (Table 1), suggesting that the experimental slower CDK2 association rate of CS242 than CS3 can be attributed to a higher energy barrier. The calculated Arrhenius parameters describing the *k*on dependence on the temperature (see Supplementary Material) indicate that hypersound irradiation increased both the frequency factor (*i.e*., from 1.3 × 10^9^ to 7.2 × 10^9^ M^-1^ s^-1^ for CS3 and from 3.9 × 10^10^ to 2.5 × 10^11^ M^-1^ s^-1^ for CS242) and the effective temperature (from 298 to 316 K (CS3) and 357 K (CS242), see Table 1). The increase in the frequency factor could be attributed to enhanced ligand diffusion (Table 1) (*16*), which would increase the collision frequency of the ligand with the protein pocket. In addition, the generation of local high-energy/pressure regions in the solvent leads to an increase in the effective temperature (*14*); however, the “macroscopic” temperature of the system remained unchanged at 298 K (Table S1). As shown in Fig. S5, the native interactions in the CDK2 structure were maintained during the hypersound-perturbed simulations, confirming that the local high-energy regions do not induce thermal denaturation of CDK2. These results suggest that the hypersound perturbation accelerates the protein-ligand association process by enhancing the cooperative local motions of the solvent molecules without affecting the native structure of the biomolecules, highlighting the general applicability of this approach for the acceleration of molecular processes in solution.

The identification of previously undiscovered druggable pockets on the protein surface plays a key role in expanding the therapeutic target range of the protein (*20*). The above conventional and hypersound-perturbed binding simulations allowed us to explore the properties of other sites beyond the ATP-binding pocket on the CDK2 surface, based on the fact that nonspecific sites (diffusion-limited) are captured by both conventional and perturbed MD, while specific (less accessible) binding sites are detected only in the hypersound-perturbed MD trajectories. We analyzed the binding simulation data of the ATP-competitive inhibitors CS3 and CS242 and two allosteric inhibitors, 2AN and 9YZ, whose binding sites are distinct from the ATP-binding pocket (*21*) (*22*). Hypersound irradiation accelerated the binding of all ligands to both the ATP-binding site (Fig. 3, green box) and the two allosteric sites 1 (Fig. 3, brown box) and 2 (Fig. 3, yellow box). Allosteric site 2 was frequently accessed by all ligands even in the conventional MD simulations (Table S3), suggesting that this site is remarkably nonspecific because of its shallow pocket shape (*22*). In contrast, the CS3/CS242 and 2AN ligands showed a more specific preference for the ATP-binding pocket and allosteric site 1, respectively, compared to their nonspecific association with allosteric site 2 (Table S3), supporting the suggestion that these ligands prefer to associate with individual binding sites, as observed in their cocrystal structures (*15, 21*). The analysis of specific and nonspecific sites based on hypersound-perturbed simulations may thus be useful for the prediction of ligand-dependent binding site selectivity and for the exploration of novel druggable cryptic sites that can allosterically regulate enzymatic activity (*20*).

**Figure 3.**
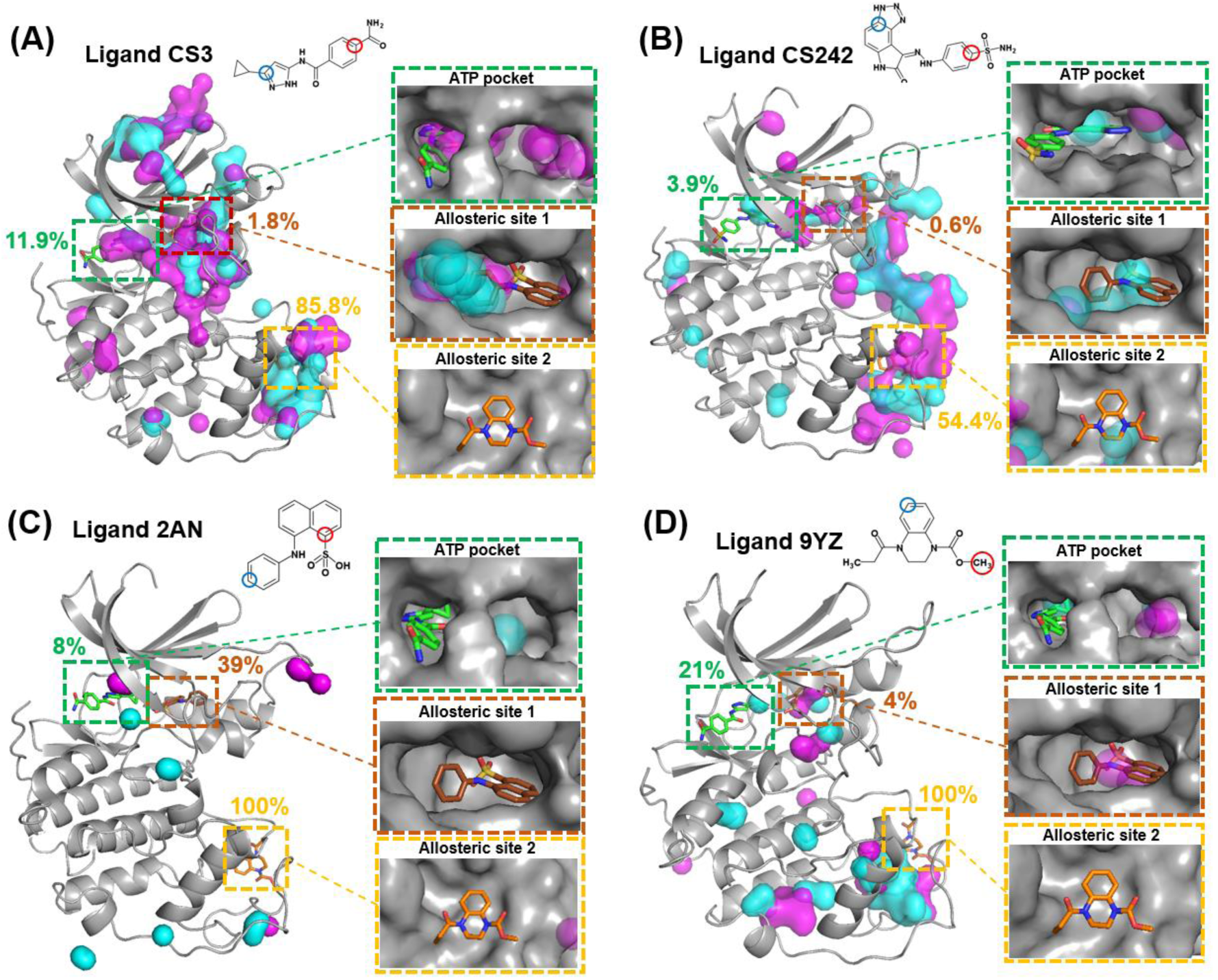
Specific binding sites of (A) CS3, (B) CS242, (C) 2AN, and (D) 9YZ ligands on the CDK2 surface predicted by comparing conventional and hypersound-perturbed MD simulations. Two representative hydrophobic and hydrophilic ligand atoms (indicated by cyan and magenta circles, respectively, in the chemical structures shown at the top of the figure) were selected for each ligand. Stable binding sites detected only in the hypersound-perturbed MD trajectories were represented in the models as semitransparent surfaces of the same color as the selected carbon atom (further details are reported in the Supplementary Material). The crystallographic poses of CS3/CS242 (green), 2AN (brown) and 9YZ (orange) inhibitors are represented as sticks. Protein structures are represented as gray ribbons or surfaces. The percentages shown in the models indicate the probabilities of capturing the ligand binding event in the hypersound-perturbed simulations (see also Table S3).

This study shows that hypersound-stimulated MD simulations have the potential to accelerate protein-ligand binding processes through a solvent-mediated mechanism without collapse of the protein structure, thus enabling the atomic-scale observation of these processes within time scales accessible by standard MD (∼ 100 ns). In contrast to other accelerated MD approaches (*23, 24*), this method does not require prior knowledge of the protein-ligand complex structure. In this way, the simulations can provide significant insights into fundamental biological mechanisms (such as the discovery of microscopic ligand binding pathways involving various bound conformations) and facilitate drug discovery (as illustrated by the present exploration of druggable binding sites on the protein surface). The present acceleration method code is publicly available (see Code Availability), can be implemented on standard high-performance computers, and is suitable for parallel computing because performing multiple independent simulations in parallel enables the sampling of a higher number of binding events and thus produces an improved statistical description of the process under study. Furthermore, the hypersound irradiation modeled in the simulations is not a fictitious computational procedure but a real physical process, even though hypersound waves of molecular (several nanometers) wavelength have not yet been realized (*16*), which currently hampers the experimental assessment of its impact. Further applications of the present technique to model other biomolecular (*e.g.*, protein conformational changes and protein-protein interactions) and non-biomolecular (*e.g*., phase transitions of materials) processes are required to assess its general effectiveness in modeling slow dynamic events.

## Abbreviations

MD: molecular dynamics
RMSF: root-mean-square fluctuation
RMSD: root-mean-square deviation
MSM: Markov state model
CDK2: cyclin-dependent kinase
PCA: principal component analysis.

## Code Availability

The hypersound-perturbed MD code is available free of charge at https://github.com/clinfo/gromacs

## Acknowledgments

This study was supported by the Ministry of Education, Culture, Sports, Science and Technology (MEXT, Japan) project “Priority Issue on Post-K Computer (Building Innovative Drug Discovery Infrastructure through Functional Control of Biomolecular Systems)”, the Foundation for Computational Science (FOCUS) Establishing Supercomputing Center of Excellence, and a Japan Society for the Promotion of Science (JSPS) KAKENHI Grant (No. JP18K06594) to MA. We thank J. Higo and I. Fukuda for critical reading of the manuscript. The reported simulations were carried out on the K computer and HPCI systems provided by the RIKEN Advanced Institute for Computational Science and Cybermedia Center, Osaka University, through the HPCI System Research Project (project IDs: hp140042, hp150025, hp150272, hp160213, hp170275, hp180186, hp190154, and ra000018).

