## Supplementary Material for "Exploring ligand binding pathways on proteins using hypersound–accelerated molecular dynamics"

**Figure S1**

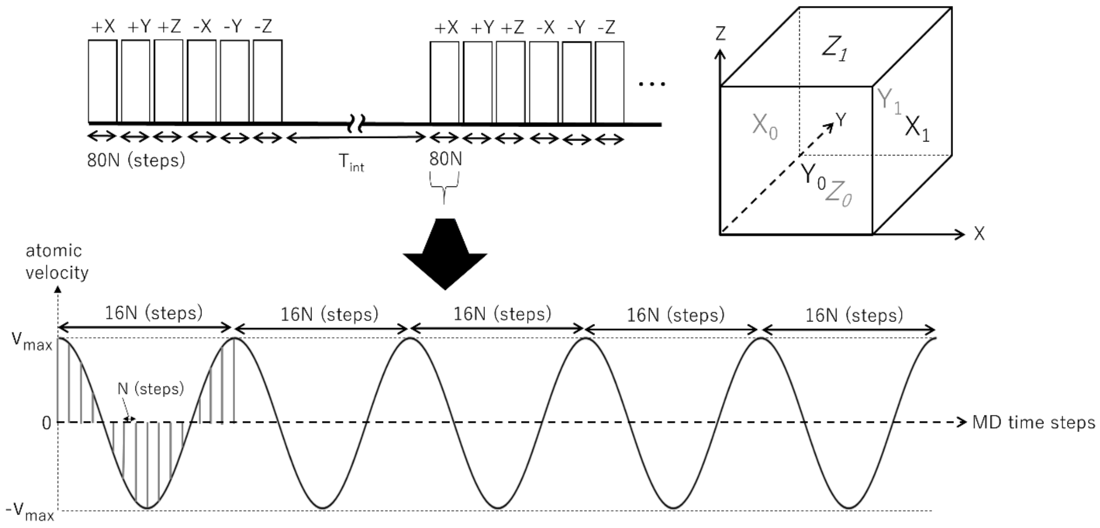


Figure S1. Schematic illustration of the modeling of shock waves in MD simulations. (top) Generation of hypersound shock waves in six directions (+X, +Y, +Z, -X, -Y, and -Z), propagating from the $X_0$, $Y_0$, $Z_0$, $X_1$, $Y_1$, and $Z_1$ surfaces, respectively, of the simulation box. (bottom) Each shock wave consisted of 5 cycles and involved 80 (=16×5 cycles) velocity pulses (indicated by vertical bars) applied every *N* MD time steps.

**Figure S2**

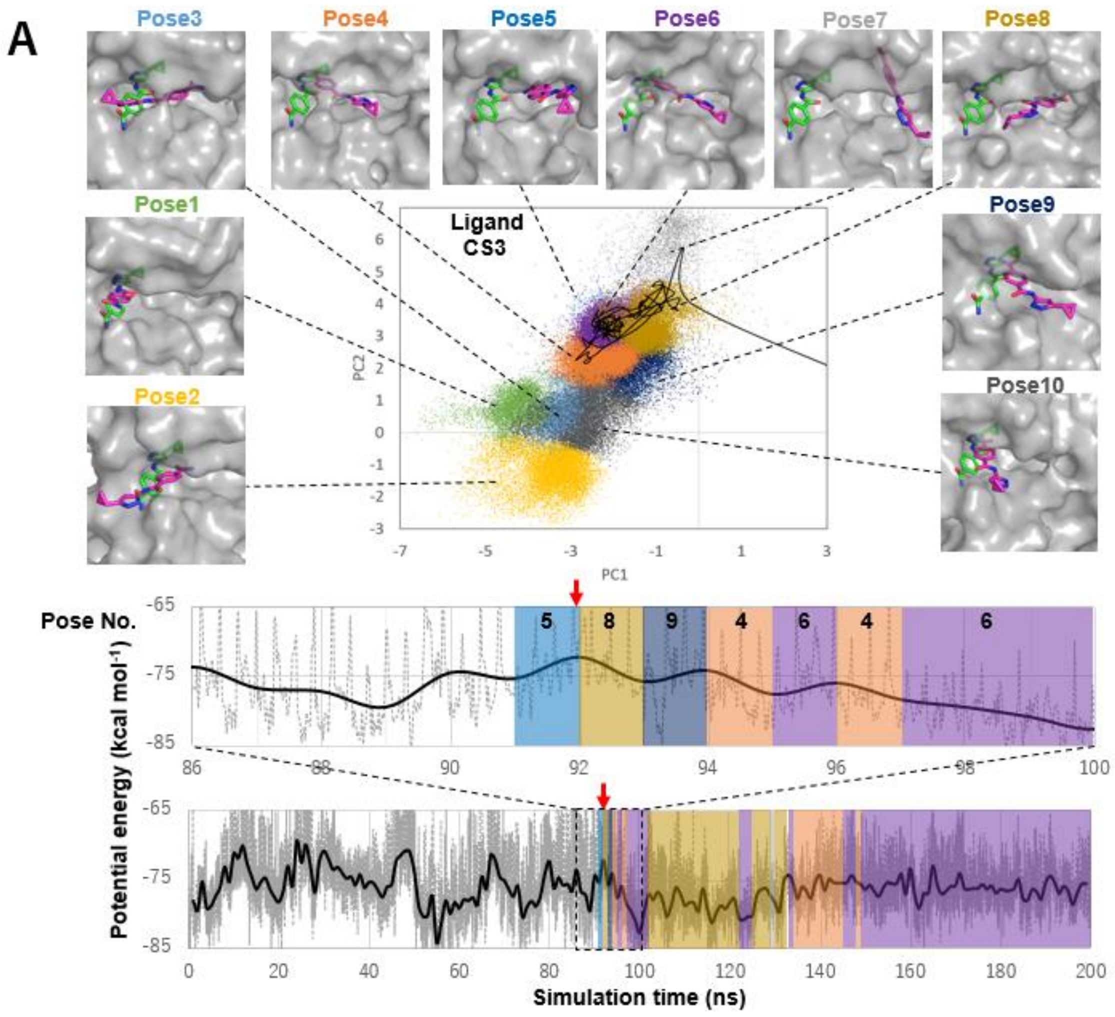
A
Pose3
Pose4
Pose5
Pose6
Pose7
Pose8
Pose1
Pose9
Pose2
Pose10
Ligand CS3
PC2
PC1
Pose No.
5
8
9
4
6
4
6
Potential energy (kcal mol-1)
Simulation time (ns)

**Figure S2 (Continued)**

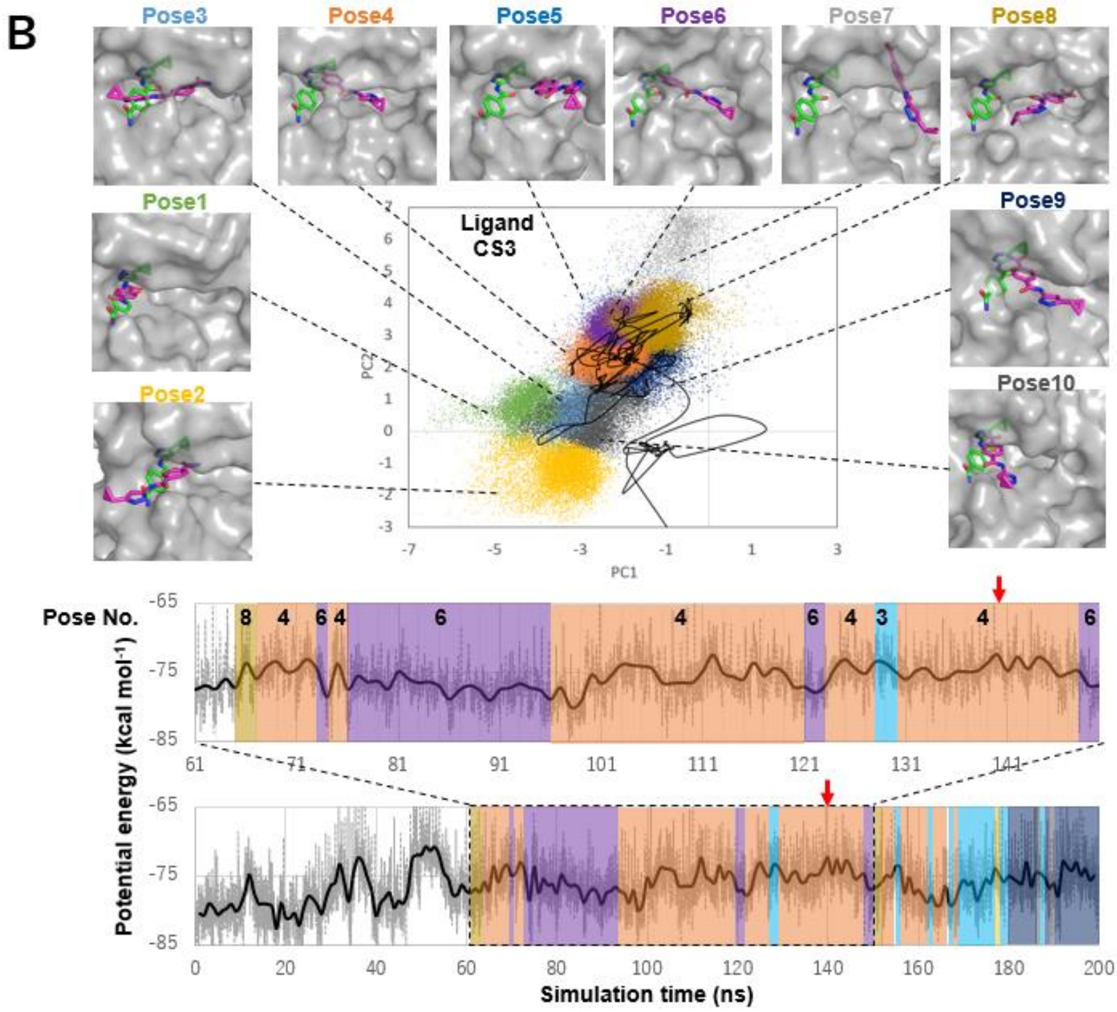

**Figure S2 (Continued)**

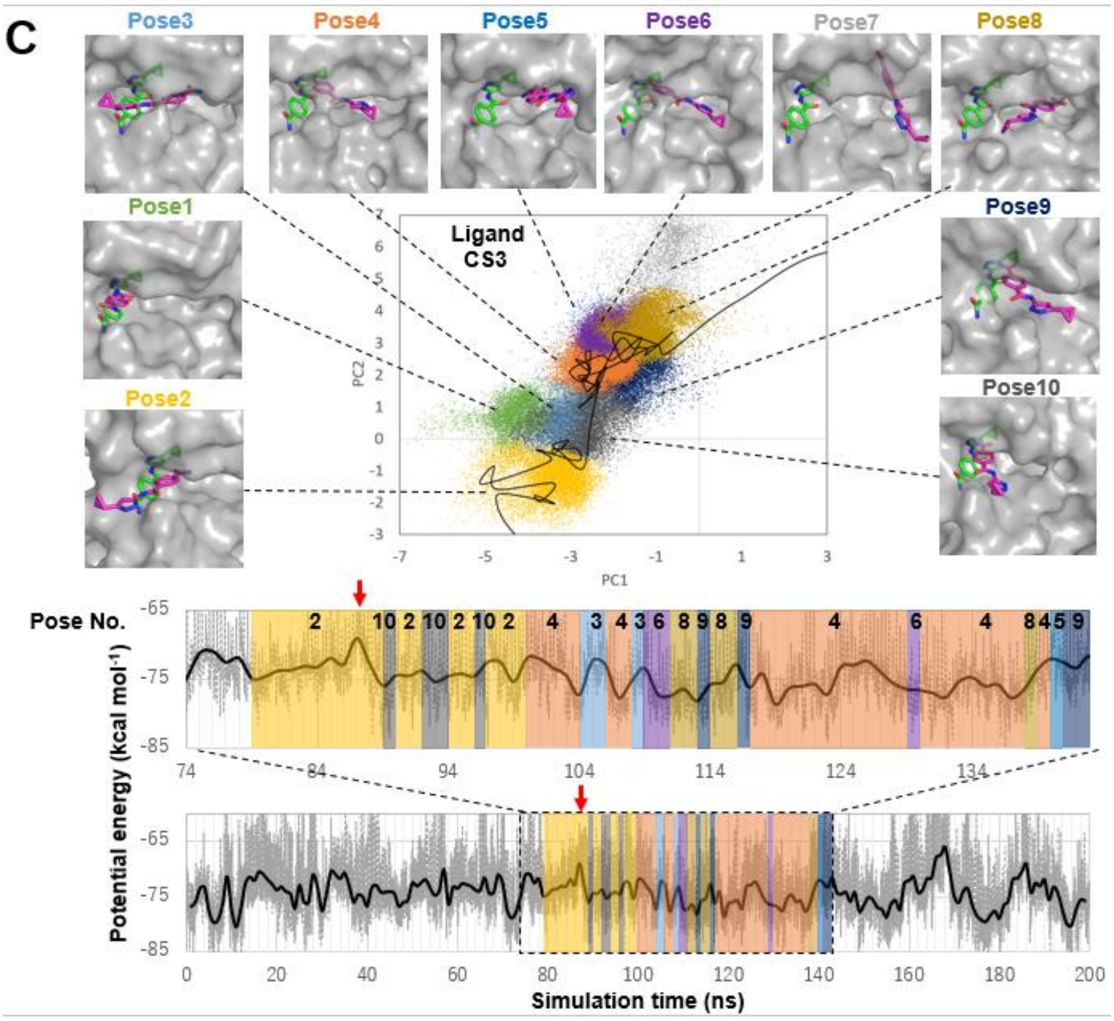

Figure S2. Three representative binding pathways of the CS3 ligand to the ATP-binding pocket of CDK2. In pathway (A), the transition state occurs upon entry into the CDK2 pocket; in (B), the transition state is reached during conformational rearrangement in the pocket interior, whereas both ligand binding and unbinding are observed in pathway (C). (top) Projections of binding conformations observed in the whole set of MD trajectories (colored dots) and of a representative binding pathway (black line) onto the first and second principal components (PC1 and PC2) calculated from PCA (see Materials and Methods section). Ten representative binding poses (magenta sticks) on CDK2 (gray surfaces) are shown along with the crystallographic pose (green sticks), the closest conformation to which was designated as Pose 1. (bottom) Potential energy trajectory corresponding to the pathway shown in the PCA map. The potential energy was calculated as the sum of the intraligand and intermolecular (protein-ligand and ligand-solvent) contributions. The highest-energy transition state is indicated by a red arrow. An enlarged view of the potential energy trajectory close to the highest-energy transition state is also shown in the panel above the whole trajectory. Time intervals in which ligand binding was observed are highlighted in the same color as that used for the binding conformation in the top panel.

### Figure S3

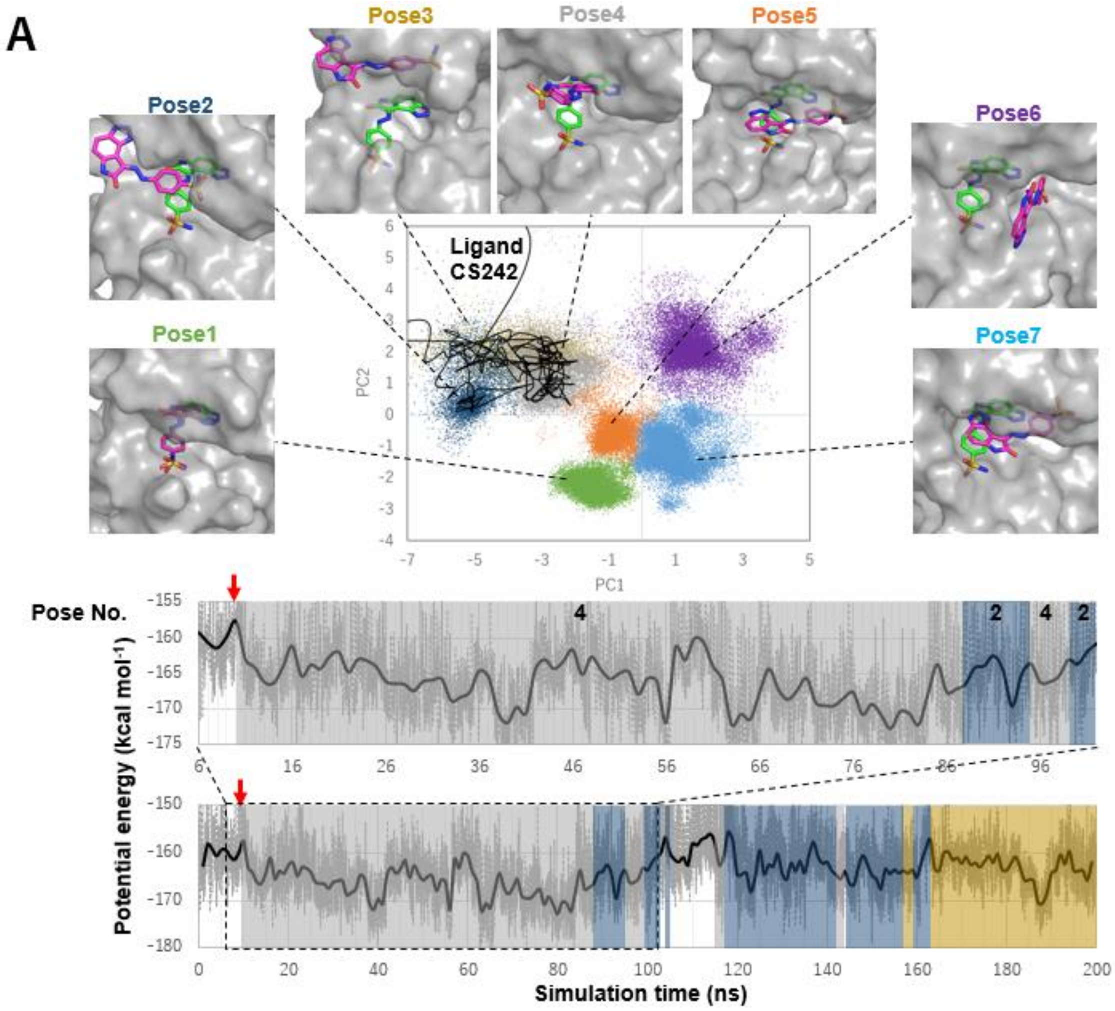
A
Pose1
Pose2
Pose3
Pose4
Pose5
Pose6
Pose7
Ligand CS242
PC1
PC2
Pose No.
4
2
4
2
Potential energy (kcal mol-1)
Simulation time (ns)

**Figure S3 (Continued)**

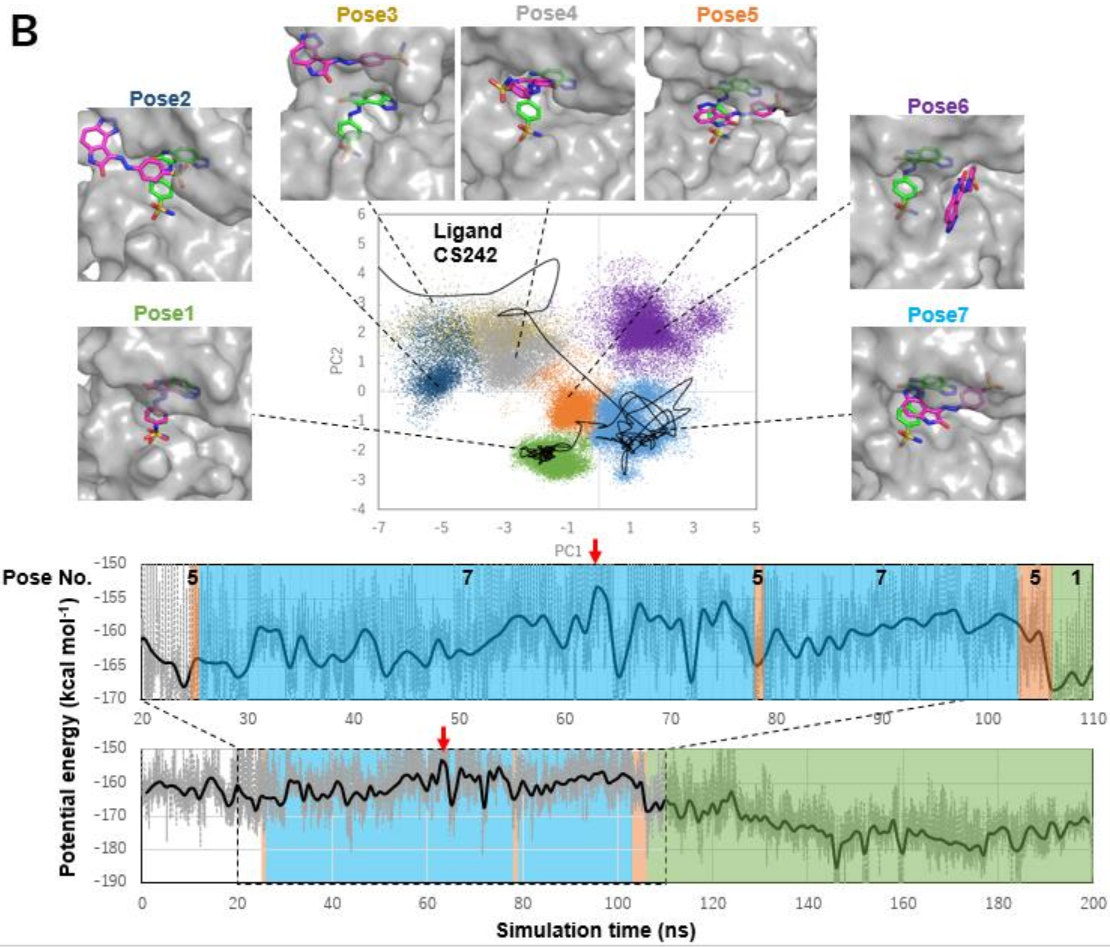

**Figure S3 (Continued)**

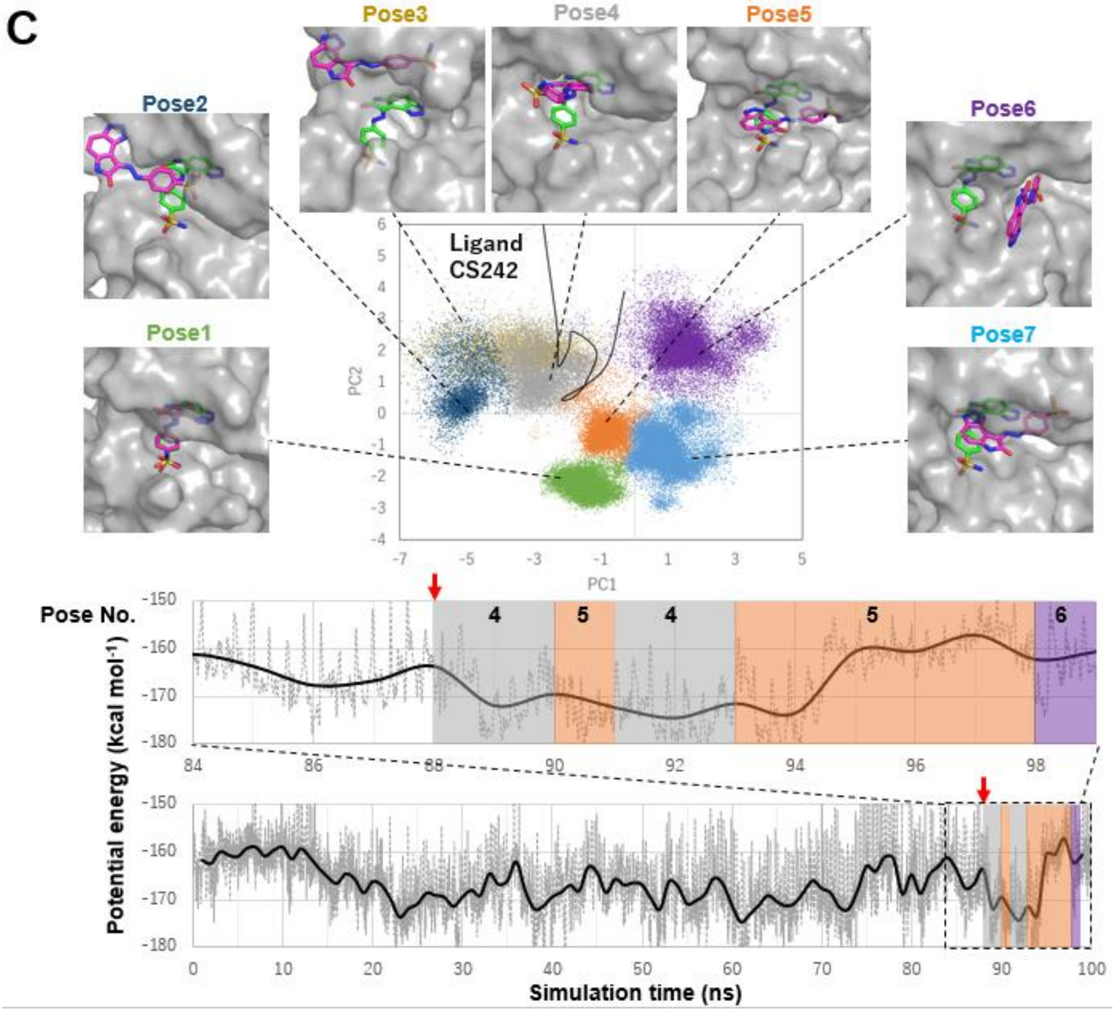

Figure S3. Three representative binding pathways of the CS242 ligand to the ATP-binding pocket of CDK2. In pathway (A), the transition state occurs upon entry into the CDK2 pocket; in (B), the transition state is reached during conformational rearrangement in the pocket interior, whereas both ligand binding and unbinding are observed in pathway (C). (top) Projections of binding conformations observed in the whole set of MD trajectories (colored dots) and of a representative binding pathway (black line) onto the first and second principal components (PC1 and PC2) calculated from PCA (see Materials and Methods section). Seven representative binding poses (magenta sticks) on CDK2 (gray surfaces) are shown along with the crystallographic pose (green sticks), the closest conformation to which was designated as Pose 1. (bottom) Potential energy trajectory corresponding to the pathway shown in the PCA map. The potential energy was calculated as the sum of the intraligand and intermolecular (protein-ligand and ligand-solvent) contributions. The highest-energy transition state is indicated by a red arrow. An enlarged view of the potential energy trajectory close to the highest-energy transition state is also shown in the panel above the whole trajectory. Time intervals in which ligand binding was observed are highlighted in the same color as that used for the binding conformation in the top panel.

**Figure S4**

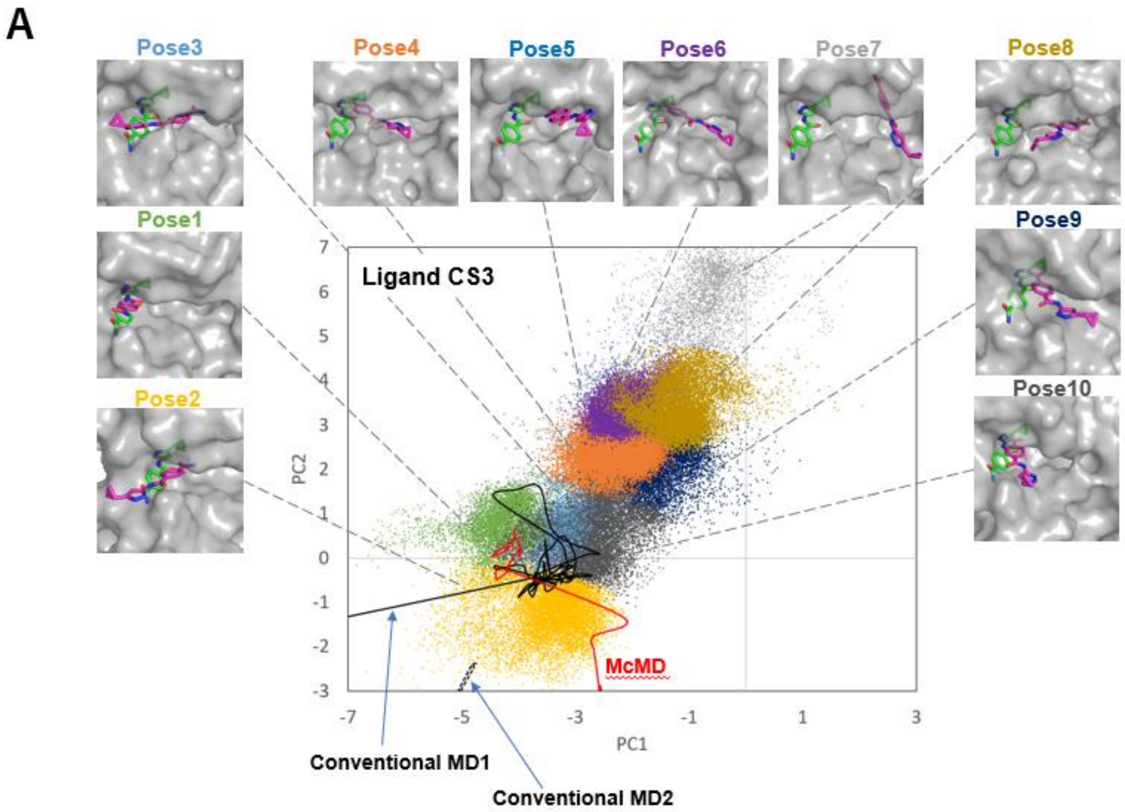
A
Pose3
Pose4
Pose5
Pose6
Pose7
Pose8
Pose1
Pose9
Pose2
Pose10
Ligand CS3
McMD
Conventional MD1
Conventional MD2
PC1
PC2

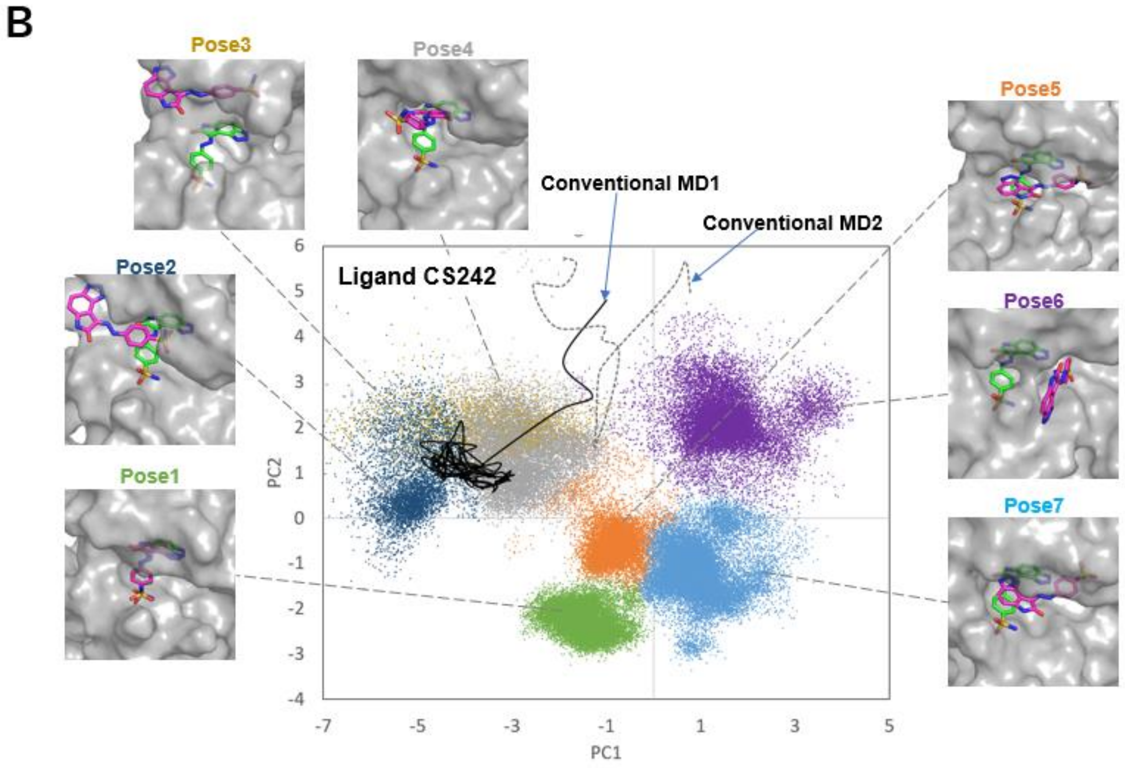
B
Pose3
Pose4
Pose5
Pose2
Pose6
Pose1
Pose7
Conventional MD1
Conventional MD2
Ligand CS242
PC1
PC2

Figure S4. Binding pathways of the CS3 (A) and CS242 (B) ligands to the ATP-binding pocket of CDK2 captured by conventional and multicanonical MD (McMD, another type of advanced MD simulations used to efficiently explore protein conformational space (*1*). Projections of binding conformations observed in the whole set of simulations (colored dots) and of representative binding pathways (lines) projected onto the first and second principal components (PC1 and PC2) calculated from PCA (see Materials and Methods section). Two binding pathways (conventional MD1 and MD2) observed in conventional MD simulations are indicated by black solid and dotted lines, while a CS3 binding pathway predicted by McMD (*2*) is indicated by a red line. Ten (CS3) or seven (CS242) representative binding poses (magenta) are shown along with the crystallographic pose (green), the closest conformation to which was designated as Pose 1.

**Figure S5**

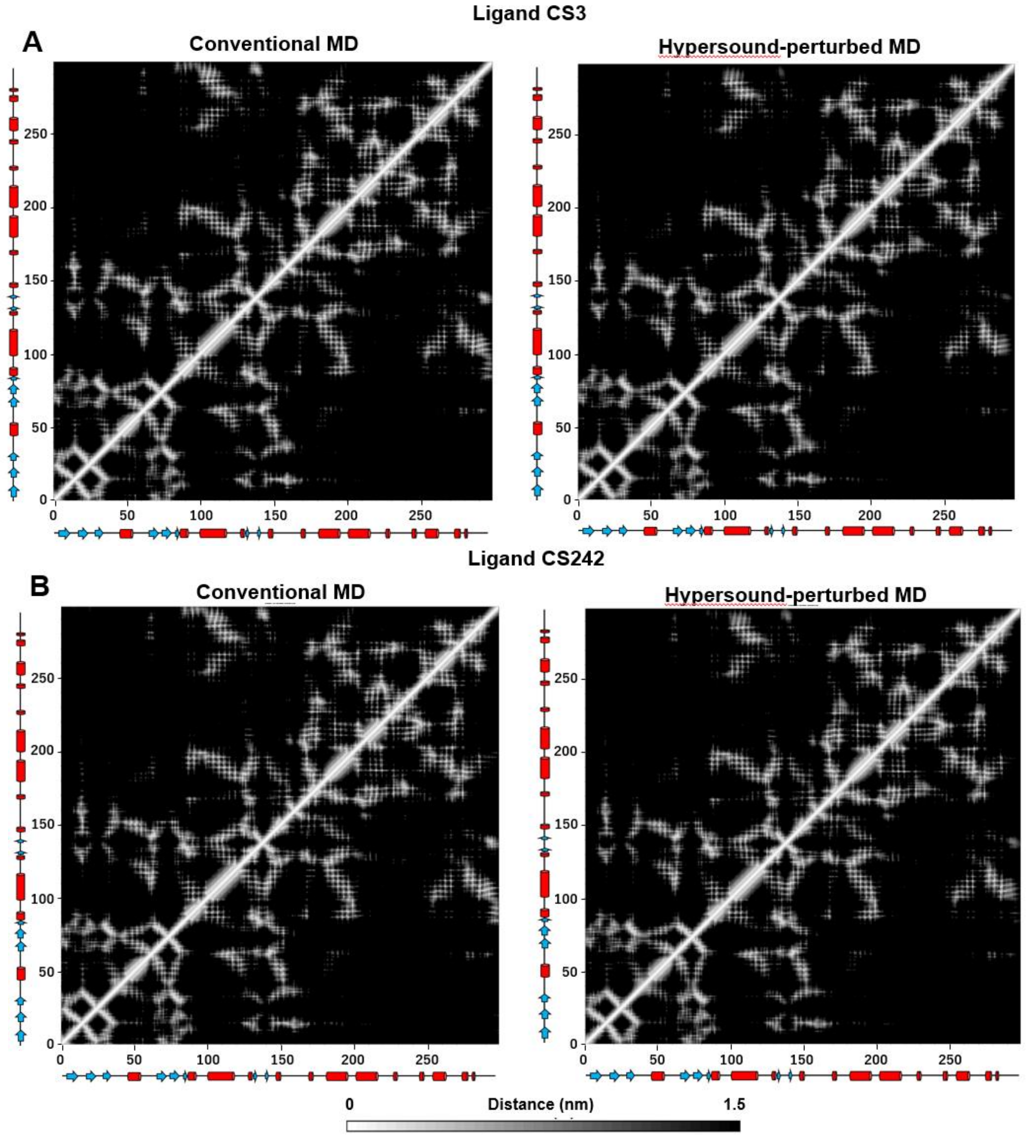


Figure S5. Native contact maps of CDK2 in the presence of (A) CS3 and (B) CS242, obtained from conventional and hypersound-perturbed MD simulations. The shortest distances between residue pairs were determined using trajectories of 50–100 ns extracted from 10 independent MD simulations of 100 ns without (left) or with (right) hypersound irradiation; no significant differences are observed in the native interaction patterns of the CDK2 structure corresponding to conventional and hypersound-perturbed MD simulations.

Table S1 Calculated thermodynamic properties of liquid water

| Simulation conditions | Temperature (K) | Diffusion constant (×10$^{-5}$ cm$^2$ s$^{-1}$) | Kinetic energy (kcal mol$^{-1}$) | Potential energy (kcal mol$^{-1}$) | Total energy (kcal mol$^{-1}$) |
|---|---|---|---|---|---|
| Conventional MD at 298 K | 298.0 ± 1.2 | 5.79 ± 0.02 | 1.78 ± 0.01 | -9.61 ± 0.01 | -7.83 ± 0.01 |
| Hypersound-perturbed MD at 298 K [a] | 295.5 ± 0.1 (297.5 ± 2.2) | 6.30 ± 0.10 (5.97 ± 0.06) | 1.76 ± 0.00 (1.78 ± 0.01) | -9.77 ± 0.01 (-9.64 ± 0.07) | -8.00 ± 0.01 (-7.86 ± 0.08) |
| Conventional MD at 305 K | 305.0 ± 1.3 | 6.40 ± 0.07 | 1.82 ± 0.01 | -9.53 ± 0.01 | -7.71 ± 0.01 |
| Conventional MD at 313 K | 313.0 ± 0.1 | 7.00 ± 0.02 | 1.87 ± 0.01 | -9.44 ± 0.01 | -7.57 ± 0.01 |
| Conventional MD at 328 K | 328.0 ± 1.3 | 8.19 ± 0.02 | 1.96 ± 0.01 | -9.28 ± 0.01 | -7.33 ± 0.01 |

[a] The thermodynamic parameters were calculated using the 0–0.048, 0.288–0.336, 0.576–0.624, 0.864–0.912, 1.152–1.200, 1.440–1.488, 1.728–1.776, 2.016–2.064, 2.304–2.352, 2.592–2.640, 2.880–2.928, 3.168–3.216, 3.456–3.504, 3.744–3.792, 4.032–4.080, 4.320–4.368, 4.608–4.656, and 4.896–4.944 ns trajectories (corresponding to the intervals in which hypersound shock waves were generated), extracted from a 5-ns MD simulation. Averages calculated over the whole 5-ns trajectory, which includes time intervals between shock wave generation ($T_{int}$ of 240 ps in Fig. S1), are indicated in parentheses.

Table S2. Summary of simulations of ligand binding to the ATP pocket of CDK2

| ligand | method | number of 100-ns simulations[a] | $N$[b] | $v_{max}$ (m $s^{-1}$)[b] | $T_{int}$[b] | number (percentage) of binding events | number of stable binding events[c] |
|---|---|---|---|---|---|---|---|
| CS3 | Conventional MD | 283 | - | - | - | 2 (0.7%) | 1 |
| | Hypersound-perturbed MD | 177 | 50 | 400 | 2,400N | 22 | 9 |
| | | 20 | 125 | 500 | 1,440N | 3 | 2 |
| | | 29 | 125 | 500 | 2,400N | 2 | 1 |
| | | total: 226 | | | | 27 (11.9%) | 12 |
| CS242 | Conventional MD | 369 | - | - | - | 2 (0.5%) | 1 |
| | Hypersound-perturbed MD | 227 | 50 | 400 | 2,400N | 11 | 6 |
| | | 64 | 50 | 300 | 2,400N | 2 | 1 |
| | | 8 | 125 | 500 | 480N | 0 | 0 |
| | | 59 | 125 | 500 | 1,440N | 1 | 0 |
| | | 4 | 125 | 700 | 2,400N | 0 | 0 |
| | | total: 362 | | | | 14 (3.9%) | 7 |
| 2AN | Conventional MD | 100 | - | - | - | 2 (2.0%) | 0 |
| | Hypersound-perturbed MD | 100 | 50 | 300 | 2,400N | 8 (8.0%) | 1 |
| 9YZ | Conventional MD | 100 | - | - | - | 7 (7.0%) | 2 |
| | Hypersound-perturbed MD | 100 | 50 | 400 | 2,400N | 21 (21.0%) | 1 |

[a] We carried out a larger number of binding simulations for CS242 than CS3 because the probability of capturing CS242 binding was expected to be lower than that of CS3, based on their experimental $k_{on}$ values ($3.35 \times 10^5$ and $3.21 \times 10^4$ $M^{-1}$ $s^{-1}$ for CS3 and CS242, respectively). Also, since the probability for conventional MDs was expected to be lower than that for hypersound-perturbed MDs, we employed the same or a larger number of conventional than hypersound-perturbed MD simulations to increase the accuracy of the probability.

[b] The $N$, $v_{max}$, and $T_{int}$ parameters are defined in Fig. S1. $N$ values of 50 and 125 correspond to hypersound frequencies of 625 and 250 GHz, respectively.

[c] Number of MD trajectories in which the formed protein-ligand complex remained stable until the end of the simulation (100 ns).

Table S3. Probabilities of observing ligand binding within different CDK2 pockets in conventional and hypersound-perturbed MD simulations[a]

| Ligand and simulation type | ATP-binding site | Allosteric site 1 | Allosteric site 2 | Number of 100-ns simulations |
|---|---|---|---|---|
| CS3 | | | | |
| Conventional MD | 0.7% (2) | 0.0% (0) | 39.2% (111) | 283 |
| Hypersound-perturbed MD | 11.9% (26) | 1.8% (4) | 85.8% (194) | 226 |
| CS242 | | | | |
| Conventional MD | 0.5% (2) | 0.0% (0) | 32.2% (119) | 369 |
| Hypersound-perturbed MD | 3.9% (14) | 0.6% (2) | 54.4% (197) | 362 |
| 2AN | | | | |
| Conventional MD | 2.0% (2) | 36.0% (36) | 99.0% (99) | 100 |
| Hypersound-perturbed MD | 8.0% (8) | 39.0% (39) | 100.0% (100) | 100 |
| 9YZ | | | | |
| Conventional MD | 7.0% (7) | 0.0% (0) | 95.0% (95) | 100 |
| Hypersound-perturbed MD | 21.0% (21) | 4.0% (4) | 100.0% (100) | 100 |

[a] The number of binding events is noted in parentheses. The maximum number of binding events per trajectory was set to one, even if multiple events were observed.

Movie S1. Hypersound-perturbed MD simulation (50 ps) of liquid water. Hypersound shock waves were sequentially generated from each of the $X_0$, $Y_0$, $Z_0$, $X_1$, $Y_1$, and $Z_1$ surfaces every 8 ps. Water molecules are displayed as red sticks.

Movie S2. MD simulation of CDK2-CS3 binding (120 of 200 ns) under hypersound irradiation. CDK2 and CS3 are displayed as surface and stick models, respectively. The corresponding potential energy [combining intraligand and intermolecular (protein-ligand + ligand-solvent) components] trajectory is also shown. The last 80 ns of the trajectory are not included in the movie because no significant conformational changes occurred in the ligand.

Movie S3. MD simulation of CDK2-CS242 binding (120 of 200 ns) under hypersound irradiation. CDK2 and CS242 are represented as surface and stick models, respectively. The corresponding potential energy trajectory [combining intraligand and intermolecular (protein-ligand + ligand-solvent) components] is also shown. The last 80 ns of the trajectory are omitted because no significant conformational changes occurred in the ligand.

### Materials and methods

#### Model systems and force fields

We modeled the binding of CDK2 to two ATP-competitive inhibitors, CS3 and CS242, and two allosteric inhibitors, 2AN and 9YZ. The initial structural data of human CDK2 were obtained from the Protein Data Bank (PDB) and the Community Structure-Activity Resource (CSAR) (http://www.csardock.org) databases (*3*). Disordered loops and flexible side chains were modeled and refined based on cocrystal structures (PDB IDs: 4EK5 (CS3), 4FKQ (CS242), 3PXF (2AN), and 5OO0 (9YZ)), and the dominant protonation state at pH 7.0 was assigned to titratable residues. Considering that a high concentration of ligands enhances the probability of capturing the protein-ligand binding (*4*), 50 ligands were randomly placed around the protein and away from the binding site (> 17 Å) by translating the ligand in the bound crystal structure.

The ligands were protonated to give net charges of 0 (CS3, CS242, and 9YZ) or -1 (2AN), reflecting the dominant protonation states at neutral pH. GAMESS was used to optimize the structure of each ligand and calculate its electrostatic potential at the HF/6-31G* level (*5*), after which the atomic partial charges were obtained *via* the restrained electrostatic potential (RESP) approach (*6*). The other potential parameters of the ligands were obtained by the general AMBER force field (GAFF) (*7*) using the antechamber module of AMBER Tools 12. The AMBER ff99SB-ILDN force field (*8*) was used for the protein and ions, while water was modeled with the TIP3P potential (*9*). Approximately 18,000 water molecules were placed around the protein model in an 8.4×8.4×8.4 $nm^3$ cubic box. In addition, approximately 60 sodium and chloride ions (corresponding to 150 mM NaCl) were introduced in the simulation box to neutralize all systems, except for the CDK2-2AN complex, for which the NaCl concentration was decreased to 10 mM because of the high concentration of the charged ligand. Based on the volume of the simulation box (592.7 $nm^3$) and the number of ligands (50), the ligand concentration was calculated to be 138 mM, which is much higher than typical concentrations used in biochemical assays, however the enhanced ligand diffusion by hypersound irradiation indicates that aggregation of ligand molecules is successfully prevented (Table 1). For the liquid water system, a total of 20,068 water molecules were included in an 8.5×8.5×8.5 $nm^3$ cubic box.

**Modeling of shock waves**

The isotropic hypersound irradiation of the solute was modeled by generating six different shock waves sequentially propagating from each face of the cubic simulation box ($X_0$, $Y_0$, $Z_0$, $X_1$, $Y_1$, and $Z_1$) toward its center (Fig. S1, top). The six shock waves were sequentially irradiated in the +X, +Y, +Z, -X, -Y and -Z directions, and a delay (corresponding to the $T_{\mathrm{int}}$ interval) was applied between each series of shock waves to prevent temperature increase. Each shock wave consisted of five cycles of 16*N* time steps: a total of 80 (= 16×5 cycles) velocity pulses were applied every *N* MD steps (Fig. S1, bottom). In each pulse, hypersound-induced velocities defined as:

$$v_i = v_{max} \times \cos(2\pi \times m / 16N) \text{ (for i = } X_0, Y_0, Z_0, \text{ m = 0, } N, 2N\ldots) \quad [1\text{-}1]$$

$$v_i = -v_{max} \times \cos(2\pi \times m / 16N) \text{ (for i = } X_1, Y_1, Z_1, \text{ m = 0, } N, 2N\ldots) \quad [1\text{-}2],$$

where $v_{max}$ is the maximum velocity assigned to the pulse and *m* is the time step number, were added to the thermal velocities of the water molecules located within 1 nm of each surface to model locally-originated shock waves. Shock waves were applied to both the liquid water and solvated protein-ligand systems. The *N*, $v_{max}$ and $T_{int}$ values used in the simulation of liquid water are reported in the next section, whereas those used in the simulations of protein-ligand binding are summarized in Table S2. A modified version of the GROMACS 4.6.5 program (*10*) was used to model the shock waves.

**MD simulations**

MD simulations with periodic boundary conditions were carried out using the GROMACS 4.6.5 program on the K computer and Cybermedia Center at Osaka University (Japan). Electrostatic interactions were calculated using the particle mesh Ewald (PME) method (*11*) with a cutoff radius of 10 Å, unless stated otherwise; van der Waals interactions were cut off at 10 Å. The P-LINCS algorithm was employed to constrain all bond lengths at their equilibrium value (*12*). After energy minimization, each system was equilibrated as described in the following subsections. A time step of 2 fs was used in all MD runs.

(i) Liquid water

The system was equilibrated for 100 ps in the constant number of molecules, volume, and temperature (NVT) ensemble. Production runs were also conducted in the NVT ensemble: the temperature was controlled with a Nose-Hoover thermostat (*13, 14*) with a time constant of 0.1 ps. Electrostatic interactions were cut off at 11 Å. Production runs of 5 ns were performed with and without hypersound irradiation. The $N$, $v_{max}$, and $T_{int}$ parameters in the hypersound-perturbed MD simulations were set to 50, 400 m $s^{-1}$, and 2,400N, respectively. The mass density, pressure, and kinetic energy of the system were calculated using the coordinates and velocities saved every 2 fs.

(ii) Protein-ligand systems

Each system was equilibrated for 100 ps under NVT conditions, followed by an MD run of 100 ps in the constant number of molecules, pressure, and temperature (NPT) ensemble, with positional restraints applied on protein heavy atoms. Production runs were then conducted under NPT conditions, without positional restraints. The temperature was maintained at 298 K by stochastic velocity rescaling (*15*), and a Parrinello-Rahman barostat was used to maintain the pressure at 1 bar (*16*). The temperature and pressure time constants were set to 0.1 and 2 ps, respectively. A total of 283, 369, 100, and 100 independent production runs of 100 ns (with different atomic velocities) were performed for the CDK2-CS3, CDK2-CS242, CDK2-2AN, and CDK2-9YZ systems, respectively. In addition, 226 (CS3), 362 (CS242), 100 (2AN), and 100 (9YZ) production runs were performed under hypersound irradiation using the parameters summarized in Table S2.

**Analysis of MD simulations of liquid water**

The mass density, pressure, and kinetic energy in the hypersound-perturbed MD simulations of liquid water were estimated by focusing on wave propagation along the X direction, as described in the following.

The mass density was estimated at 82 different X-points, based on the number of molecules located within ±0.2 nm of each point. The kinetic energy ($k_x$) was calculated as $k_x = \frac{1}{2}M\langle v_x\rangle^2$, where $M$ is the mass of a water molecule and $\langle v_x \rangle$ is the X component of the velocity, averaged over all water molecules located within ±0.2 nm from the corresponding X-point. Under hypersound irradiation, $k_x$ was estimated to be 0.4–0.5 kcal/mol at the center of the simulation box (X = 4 nm, Fig. 1C). The instantaneous temperature in this region was estimated to be 400–500 K, based on the $k_x$ value of bulk water at 300 K (~ 0.3 kcal/mol, corresponding to *RT*/2).

The pressure of water in the +X direction of the cubic simulation box was estimated from the X components of the velocities of the water molecules that crossed the YZ plane at a given X during the observation time $\Delta t$, according to the modified van der Waals equation for liquid systems:

$$P = \frac{2m}{S\Delta t}\sum_i v_x^i - a\left(\frac{N_a}{V_m}\right)^2 \quad [2]$$

where $m$ is the mass of a water molecule, $S$ is the area of the YZ plane, $v_x^i$ is the X component of the velocity of the $i$-th water molecule, $a$ is the intermolecular attractive force constant (determined as described below), $N_a$ is the Avogadro's number, and $V_m$ is the molar volume, which was calculated to be 0.0183 L/mol based on the volume of the simulation box ($8.5^3$ $nm^3$) and the number of water molecules contained in it (20,068). We initially performed a conventional MD simulation of 50 ps, and the first term of equation [2], $\frac{2m}{S\Delta t}\sum_i v_x^i$, was calculated to be $1.298 \times 10^8$ Pa based on the water molecules that crossed the YZ plane at X = 2 nm (corresponding to the mid-point between the origin and the center of the simulation box) during a $\Delta t$ interval of 50 ps. Using the saturated vapor pressure of water at 298 K ($P = 0.032 \times 10^5$ Pa), the $a$ parameter was then estimated to be 0.423 (atm $L^2$ $mol^{-2}$). The pressure under hypersound irradiation was then determined from the hypersound-perturbed MD trajectory, using the estimated $a$ value and the sum of the $v_x$ values of the water molecules that crossed the YZ plane at each selected X point during a $\Delta t$ interval of 0.4 ps.

**Analysis of ligand binding within different CDK2 pockets**

For each ligand, we analyzed the MD trajectories of the system containing the CDK2 protein and 50 ligand molecules. Ligand binding within individual CDK2 sites (ATP pocket, allosteric site 1, and allosteric site 2) was considered to occur if at least two distances between an atom belonging to the protein pocket (see below) and any ligand atom were below 5 Å. The following atoms of the protein pocket were used in the distance calculation: Val18 (beta carbon, Cβ) and Leu134 (gamma carbon, Cγ) for the ATP pocket, Tyr15 (zeta carbon, Cζ) and Leu55 (gamma carbon, Cγ) for the allosteric site 1, and Cys177 (gamma carbon, Cγ) and Trp227 (indole nitrogen, Nε) for the allosteric site 2.

**Advanced analysis of CS3 and CS242 binding to the ATP pocket of CDK2**

For the ATP-competitive inhibitors (CS3 and CS242), whose experimental binding structures and $k_{on}$ values are available, the occurrence of a binding event to the ATP pocket was assessed by a stricter criterion, as follows. First, we identified trajectories that satisfied two conditions: (1) a distance between the Val18 Cβ and any ligand atom ≦ 5 Å and (2) the RMSD of the ligand from the crystallographic pose below 9 Å. Next, entry into the ATP pocket was confirmed by visual inspection of these trajectories, using the VMD software (*17*). Finally, we identified 27 (CS3) and 14 (CS242) MD trajectories that captured binding events.

In approximately half of these MD trajectories, the bound state was unstable, and the ligand separated from the ATP pocket within 1–40 ns. However, in the remaining trajectories, the ligand remained stably bound to the protein until the end of the simulation; these trajectories (12/27 for CS3, 7/14 for CS242) were thus extended to 200 ns to further examine the behavior of the bound ligands.

Principal component and conformational clustering analyses of the ligand binding poses observed in the 27 (CS3) and 14 (CS242) MD trajectories were performed as follows: after removing the overall translation and rotation of the protein, the covariance matrix was calculated using the Cartesian coordinates of the ligand and diagonalized to obtain the PC eigenvectors. Conformational clustering of the binding poses into an optimal number of clusters was then performed on the first three PCs (PC1–PC3) using the X-means clustering method (*18*). The bound states of CS3 and CS242 on the CDK2 surface were grouped into 10 and 7 conformational clusters, respectively, one of which corresponded to the crystallographic pose (*3*), indicating that some of these binding conformations are commonly observed in the 27 (CS3) and 14 (CS242) trajectories.

**Estimation of kinetic parameters for the CDK2-ligand binding process**

The association rate constant under hypersound irradiation ($k_{on}$), activation energy ($E$), diffusion constant of the solute ($D$), steric factor ($P$), frequency factor ($A$), and effective temperature under hypersound irradiation ($T$) were estimated as follows, using the experimental $k_{on}$ values measured without any perturbation and the trajectories obtained from conventional and hypersound-perturbed MD simulations.

The kinetics of the binding between protein (P) and ligand (L) was analyzed according to the following reaction scheme:

$$\mathrm{P} + \mathrm{L} \xrightarrow{k_{on}} \mathrm{PL}$$

where PL is the protein-ligand complex.

The second-order reaction rate is defined as:

$$\frac{d[\mathrm{PL}]}{dt} = k_{on}[\mathrm{P}][\mathrm{L}] \qquad [3]$$

where [P], [L], and [PL] are the concentrations of the protein, ligand, and protein-ligand complex, respectively. The initial binding rate is proportional to the initial concentrations of P and L ($[\mathrm{P}]_0$ and $[\mathrm{L}]_0$, respectively). If $[\mathrm{P}] \approx [\mathrm{P}]_0$ and $[\mathrm{L}] \approx [\mathrm{L}]_0$, the following relation can be derived by solving equation [3]:

$$\frac{[\mathrm{PL}]}{[\mathrm{P}]_0} = k_{on}[\mathrm{L}]_0 t \qquad [4]$$

The $[\mathrm{P}]_0$ and $[\mathrm{L}]_0$ values in the present simulations of the CDK2-ligand binding were 2.8 and 138 mM, respectively. Based on the experimentally determined $k_{on}$ values of CS3 and CS242 [$3.35 \times 10^5$ and $3.21 \times 10^4$ $M^{-1}$ $s^{-1}$, respectively (*3*)], the fractions of the CDK2-ligand complexes after 100 ns were expected to be 0.46% (CS3) and 0.044% (CS242). The probabilities of observing the stable ligand binding event in the 100-ns conventional MD simulations of CS3 and CS242 were 0.4% (= 1/283) and 0.3% (= 1/369) (Table S2), respectively. Under hypersound irradiation with $N$ = 50 steps, $v_{max}$ = 400 m/s, and $T_{int}$ = 2,400 N, which are the parameters predominantly used in the simulations (Table S2), and using $[\mathrm{PL}]/[\mathrm{P}]_0$ ratios of 9/177 (CS3) or 6/227 (CS242), corresponding to the proportions of MD trajectories that exhibited stable ligand binding (Table S2), the $k_{on}$ values were estimated to be $3.68 \times 10^6$ (CS3) and $1.92 \times 10^6$ $M^{-1}$ $s^{-1}$ (CS242).

The $k_{on}$ constant can also be described using the Arrhenius equation:

$$k_{on} = Aexp\{-\frac{E}{RT}\} \quad [5]$$

where $R$ is the gas constant. To estimate $E$, the potential energy difference between the unbound state and the highest-energy transition state was averaged over 12 (CS3) and 7 (CS242) trajectories, corresponding to all hypersound-perturbed MD trajectories that exhibited stable ligand binding (Table S2). According to a kinetic model involving a "doorway state" located between the unbound and bound states, the frequency factor can be approximated by the diffusion-controlled rate constant (*19*):

$$A = 4\pi N_A(D_P + D_L)R^*P \quad [6]$$

where $N_A$, $D_P$ ($D_L$), $R^*$, and $P$ are the Avogadro's number, the diffusion constant of the protein (ligand), the critical protein-ligand distance, and a steric factor, respectively. In this study, $R^*$ was set to $10^{-9}$ nm, and we assumed $D_P << D_L$. To calculate $D_L$, the mean-square displacement of each ligand during an MD simulation was averaged over 50 independent simulations. The diffusion constants ($D_{L\_conv}$) of CS3 and CS242 estimated from the conventional MD simulations were $1.87 \pm 1.33 \times 10^{-5}$ and $1.55 \pm 1.01 \times 10^{-5}$ $cm^2/s$, respectively, while those estimated from the MD runs under hypersound irradiation with $N$ = 50 steps and $v_{\max}$ = 400 m/s ($D_{L\_hyper}$) were $10.62 \pm 6.98 \times 10^{-5}$ $cm^2/s$ (CS3) and $9.87 \pm 7.00 \times 10^{-5}$ $cm^2/s$ (CS242). Using the $D_{L\_conv}$ and experimental $k_{on}$ values along with the $E$ parameter in equations (5) and (6), the steric factors of CS3 and CS242 were calculated as $10^{-1.05\pm1.64}$ and $10^{0.52\pm1.70}$, respectively. According to equation (6), the frequency factors ($A$) without hypersound irradiation were calculated to be $10^{9.10\pm1.69}$ (CS3) and $10^{10.59\pm1.73}$ (CS242), while those obtained under hypersound irradiation were $10^{9.86\pm1.68}$ (CS3) and $10^{11.39\pm1.74}$ (CS242). Finally, the effective temperatures under hypersound irradiation calculated from equation (5) were 316 K for CS3 and 357 K for CS242.

**Identification of specific ligand binding sites on the CDK2 surface**

Two representative carbon atoms belonging to the hydrophobic and hydrophilic groups were selected for each ligand (Fig. 3). First, the root-mean-square fluctuation (RMSF) of each atom was calculated every 10 ns of the individual 100-ns conventional MD trajectories, and if the value was below 3 Å, the mean coordinates of the atom were recorded as nonspecific (diffusion-limited) binding site. Next, the same protocol was applied to the hypersound-perturbed MD trajectories, and binding sites detected only in the latter were identified as specific (less accessible) sites.
